# Metabolic evolution in response to interspecific competition in a eukaryote

**DOI:** 10.1101/2022.10.19.512836

**Authors:** Giulia Ghedini, Dustin J. Marshall

## Abstract

Competition can drive rapid evolution which, in turn, alters the trajectory of ecological communities. The role of eco-evolutionary dynamics in ecological communities is increasingly well-appreciated, but a mechanistic framework for identifying the types of traits that will evolve, and their trajectories, is required. Metabolic theory makes explicit predictions about how competition should shape the evolution of metabolism and size but these predictions have gone largely untested, particularly in eukaryotes. We use experimental evolution of a eukaryotic phototroph to examine how metabolism, size, and demography coevolve under both inter- and intra-specific competition. We find that the focal species evolves a smaller body size in response to competition, reducing density-dependence and maximizing carrying capacity. Metabolic theory successfully predicted most of these adaptations, but we also find important departures from theory. Longer-term evolution (70 generations) led to Pareto improvements in both population growth rate and carrying capacity, suggesting that classic r-K trade-offs observed among species can be evaded within species. The evasion of this trade-off appeared to arise due to the rapid evolution of enhanced metabolic plasticity: lineages exposed to competition evolved more labile metabolisms that tracked resource availability more effectively than lineages that were competition-free. We predict that rapid evolution in both size and metabolism may be a ubiquitous feature of adaptation to changing resource regimes that occur *via* species invasions and environmental change.

## Introduction

When ecological and evolutionary processes occur on similar timescales, eco-evolutionary dynamics can shape the diversity and functioning of communities ^1–3^. Studies in bacteria imply that competition from whole communities alters evolutionary trajectories in ways that are fundamentally different from simpler pairwise interactions ^4–7^. Whether these results extend to eukaryotic communities, where species tend to have lower population densities and longer life cycles, and thus slower rates of evolution, remains unclear ^8^. Macroevolutionary patterns in eukaryotes suggest that competition for resources can drive species to evolve differential resource use (e.g. Darwin’s finches) ^9–11^ but such escapes from competition may not always be accessible ^12,13^. In most cases, predicting trait evolution in species that share similar niches remains a formidable challenge ^8,14,15^.

Metabolic theory should help with identifying the eco-evolutionary trajectories of key phenotypic traits. The theory proposes that metabolism should drive competition by setting *per capita* resource demands ^16,17^. For example, because smaller organisms have lower absolute metabolic rates, their populations should support higher maximum population densities than those of larger organisms ^18^. Similarly, because small organisms usually have higher metabolism per unit mass (assuming that metabolism scales hypo-allometrically with body size), their populations should grow faster than those of larger organisms – producing classic r-K trade-offs often observed among species ^17^. Therefore, evolved changes in size and metabolism should both affect demography and capacity to compete with other species ^19,20^ but whether size and metabolism actually coevolve in response to interspecific competition has not been explored.

We combined experimental evolution and metabolic theory to determine how size, metabolic fluxes, and demography coevolve in response to both intra- and inter-specific competition in phytoplankton. We base our assessment on a focal species, the marine eukaryotic microalga *Dunaliella tertiolecta*, which we evolved for ~70 generations in each of three environments: i) competition-free; ii) intraspecific competition; and iii) interspecific competition (three other phytoplankton species). To manipulate the environment, we used dialysis bags that physically isolated the evolving population from its competitors while exposing it to competition for light and nutrients ^6,21–23^ (Fig. 1). We treated the competition-free lineages as a reference control. We batch-transferred both the focal species and the competitors weekly standardizing biovolumes between treatments (~7 generations). We assessed how the focal populations had evolved at two points: at ~35 generations and ~70 generations. At each of these time points, we placed subsamples of all the evolved lineages of the focal species into a common environment, and left them for several generations such that any persistent phenotypic differences that remained among treatments represented evolved responses to the environments ^24^. We assessed metabolism (photosynthesis, respiration, and net daily energy production), morphology (cell size and shape) and demography (growth rate, max. population density, and max. total biomass).

**Figure 1.**
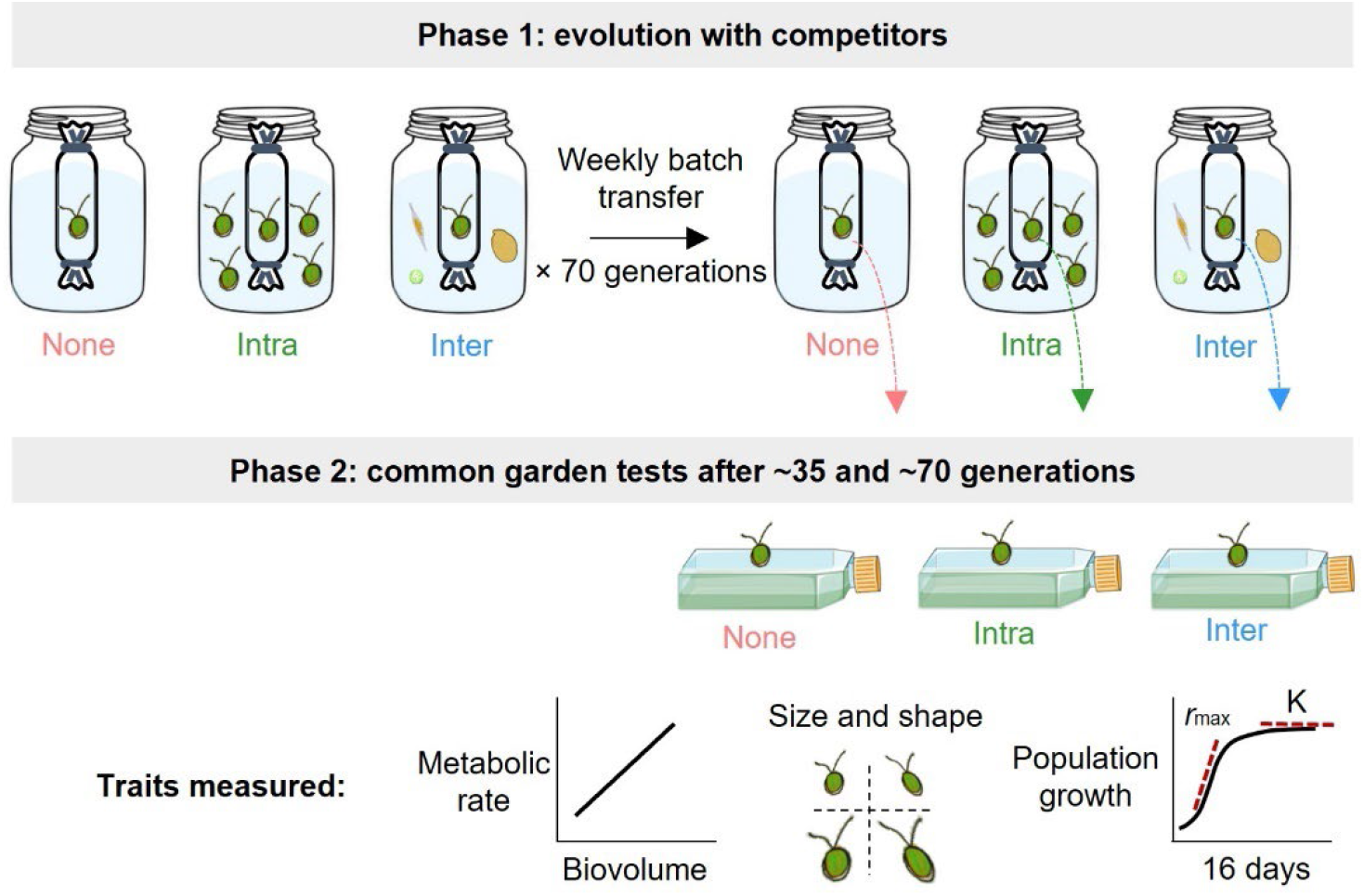
To test how metabolism, size and demography coevolve in response to competition, we evolved the eukaryotic microalga *Dunaliella tertiolecta* in three environments: competition-free (“none”), intraspecific (“intra”) and interspecific competition (“inter”). We enclosed the focal species in dialysis bags that physically isolated the evolving population from competitors while exposing it to competition for light and nutrients for 10 weeks corresponding to ~70 generations. We propagated the focal lineages and the competitors performing weekly batch transfers. After ~35 and ~70 generations we used common garden experiments to compare changes in metabolic fluxes, morphology, and demography among the evolved focal populations.

## Results

### Metabolism evolves under competition

After ~35 generations of experiencing interspecific competition, algae evolved lower metabolic fluxes than algae that experienced no interspecific competition. Specifically, rates of both photosynthesis and dark respiration were lower in lineages exposed to interspecific competition (photosynthesis: competition × biovolume interaction, F_2,270_ = 3.85, p = 0.02; respiration: main competition effect, F_2,272_ = 6.56, p = 0.002; see Supplementary Information: Fig. S1, Table S1). These differences in metabolism were only apparent when we measured energy fluxes under resource limitation; the lineages were the same when resources were abundant (Fig. S1, Table S1). In other words, algae that experienced interspecific competition evolved plasticity to have ‘thriftier’ metabolisms when resources were scarce, reducing their rates of energy use, and presumably nutrient demands, while photosynthesising relative to other lineages.

After a further 35 generations of experimental evolution, interspecific competition drove more pronounced evolutionary changes in metabolism and metabolic plasticity. When we assessed metabolic fluxes under resource-limiting conditions, metabolic rates (both photosynthesis and respiration) were lowest in lineages that had experienced interspecific competition and highest in those that experienced no competition (competition effect on photosynthesis: F_2,47_ = 6.75, P = 0.003; Fig. 2b; respiration: F_2,47_ = 15.83, p < 0.0001, Fig. 2d; Table S1). Again, lineages that evolved in the presence of competition were thriftier when resources were scarce, particularly those that evolved under interspecific competition. But when we assessed metabolic fluxes when resources were abundant, the patterns were reversed: lineages exposed to competition had the highest rates of photosynthesis and respiration relative to competition-free lineages. In other words, lineages that experienced competition had evolved greater capacity to both fuel (via photosynthesis) and do (via respiration) biological work when resources were abundant (competition × biovolume effect on photosynthesis: F_2,210_ = 3.59, p = 0.03; Fig. 2a; competition effect on respiration: F_2,46_ = 10.86, p = 0.0001, Fig. 2c; Table S1). Overall then, competition-exposed lineages evolved higher levels of metabolic plasticity than competition-free lineages: competition-exposed lineages downregulated their resource demands when resources were scarce and strongly upregulated their demands during periods of resource abundance (Fig. 3). In contrast, competition-free lineages exhibited more modest metabolic plasticity in response to changes in resource availability. Together, these evolved metabolic changes meant that competition-exposed lineages produced more energy on balance (photosynthesis – respiration), thereby fuelling more biological work when resources were abundant (Fig. 2e), but required fewer resources and produced less net energy when resource were scarce, relative to competition-free lineages (Fig. 2f).

**Figure 2.**
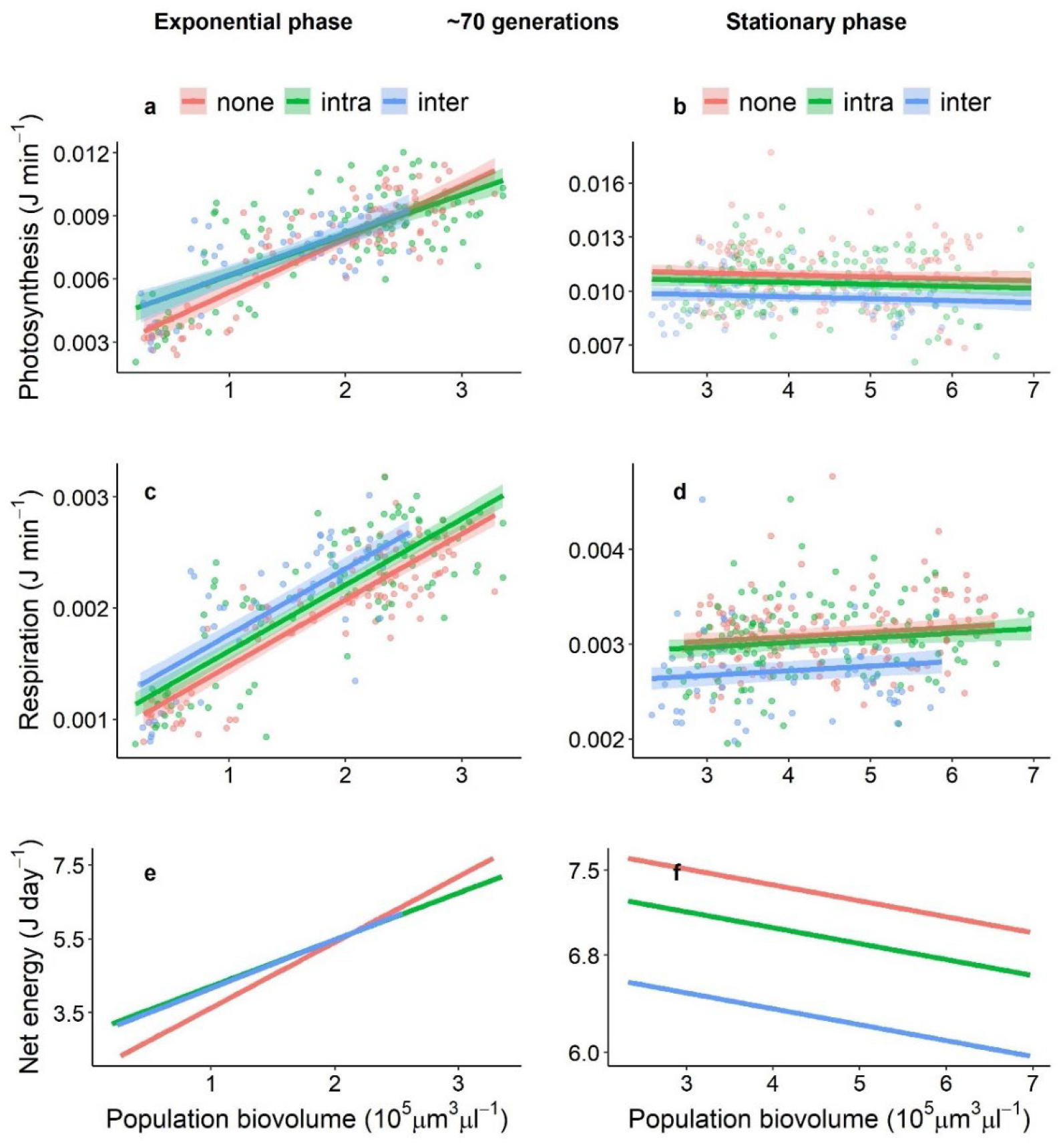
Populations exposed to interspecific competition for ~70 generations evolved greater metabolic plasticity (as revealed by the relationship between population biovolume and metabolic rate) than competition-free populations. Competition exposed populations had greater (a) photosynthesis and (c) respiration rates when resources were abundant during the exponential growth phase (left panel). But upon entering stationary phase and approaching carrying capacity (right panel), populations that experienced competition from interspecifics had much lower metabolic rates (b for photosynthesis, d for respiration). Integrating photosynthesis and respiration over 24 hours, we found that populations that had evolved in the presence of competitors produced more net energy than competition-free populations when resources were abundant (e) but they also produced less net energy when resources were scarce because they downregulated their metabolism relatively more (f).

**Figure 3.**
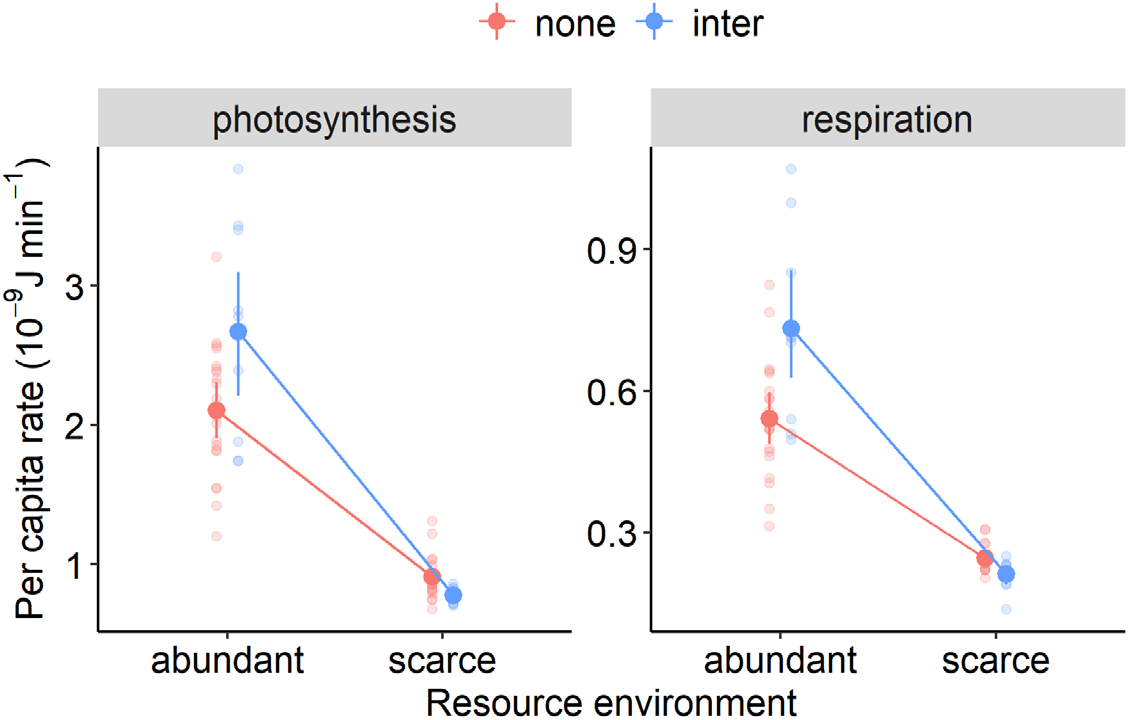
Populations exposed to interspecific competition for ~70 generations (“inter”) evolved greater metabolic plasticity relative to lineages that experienced no competition (“none”) allowing them to better track resource abundance. Cells with a history of competition increase their metabolic rates (both photosynthesis and respiration) much more than control lineages when resources are abundant (day 3, exponential growth phase) but downregulate them when resources are scarce (day 11, stationary phase).

### Size evolution, metabolic scaling and predicting demography

Lineages exposed to competition evolved smaller cell sizes than lineages that were free from competition (Fig. 4a for day 3 after 10 weeks; estimated marginal means for day 3: competition-free = 202; intra = 185; inter = 162 μm^3^). After 70 generations of experimental evolution, lineages experiencing interspecific competition were on average 13.4% smaller in cell volume than competition-free lineages (ranging from 3.6% to 19.6% smaller during common garden experiments), while those experiencing only intraspecific competition were intermediate in size (5.5% smaller on average; ranging from 0.6% larger to 8.5% smaller in size than competition-free lineages) (competition × time: F_24,564_ = 3.65, p < 0.0001; Fig. S2 for all days, Table S2). Cell shape coevolved with cell size: lineages exposed to competitors evolved rounder cells than competition-free lineages (Fig. 4b for day 3; Fig. S2; competition × time: F_24,564_ = 8.50, p < 0.0001, Table S3 for post-hoc comparisons).

**Figure 4.**
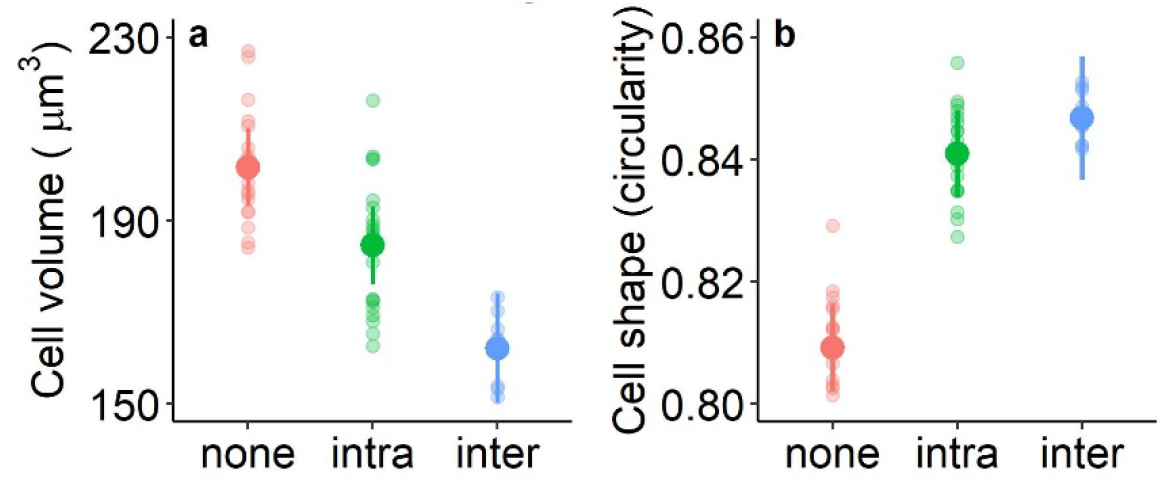
Both the size and shape of the microalga *Dunaliella tertiolecta* evolved in response to competition. After ~70 generations, populations exposed to competitors evolved smaller cell sizes, with a stronger decline in response to interspecific rather than intraspecific competitors (a, here shown for day 3). Changes in cell size were accompanied by changes in shape: the smaller cells exposed to intra- and inter-specifics competitors were also rounder than competition-free cells. See Fig. S2 for the complete temporal series.

While metabolic rates often scale hypo-allometrically for eukaryotes, we found that both photosynthesis and respiration rates scaled hyper-allometrically with cell size in the focal species (photosynthesis: 1.31 [CI: 1.05; 1.58]; respiration: 1.28 [1.02; 1.55]; Table S4), similarly to what previously found for this species ^20^. But the scaling became progressively shallower as populations grew denser over time (Table S4). Thus, under hyper-allometry, the smaller cells evolved under interspecific competition should have lower resource demands *per capita*, in an absolute sense but also relative to cell biovolume. Given the observed metabolic scalings during the exponential growth phase (*B_exponential_* = 1.13) and stationary phase (*B_stationary_* = 0.60), metabolic theory would therefore make the following predictions regarding the scaling of demographic parameters with cell size ^19,20,25^:

1. *r*_max_ = M^B^_exponential_/M^1^ = M^0.13^
2. K_cells_ = M^0^/M^B^_stationary_ = M^−0.60^
3. K_bio_ = M × K_cells_ = M × M^−B^ = M^0.40^

From these equations, we qualitatively predict that competition exposed lineages (which evolved smaller sizes) should have lower intrinsic rates of increase (*r_max_*) but higher carrying capacities (K_cells_) than competition-free lineages. In a quantitative sense, we also predict that growth rate and biomass carrying capacity (K_bio_) should positive covary with cell size, while population carrying capacity should negatively covary with cell size (K_cells_). We tested both sets of predictions experimentally.

### The evolution of greater carrying capacity

We found strong qualitative and quantitative support for our demographic predictions based on metabolic theory. After ~35 generations, lineages that experienced competition had lower maximum growth rates but greater carrying capacity (K_cells_) than competition free lineages, displaying a classic r-K trade-off (effect of competition on *r_max_*: F_2,42_ = 10.46, p < 0.0005, Fig. 5a; effect on K_cells_: F_2,42_ = 19.18, p < 0.001, Fig. 5b). After 70 generations of experimental evolution, we found the same evolved difference in carrying capacity (K_cells_; F_2,46_ = 13.91, p < 0.001, Fig. 5d) but the competition-exposed lineages showed maximum population growth rates that were similar to the competition-free lineages (F_2,46_ = 1.97, p = 0.15, Fig. 5c; Table S5). The evolved demographic responses were similar regardless of whether competition was intra- or inter-specific (Fig. S3, Table S5).

**Figure 5.**
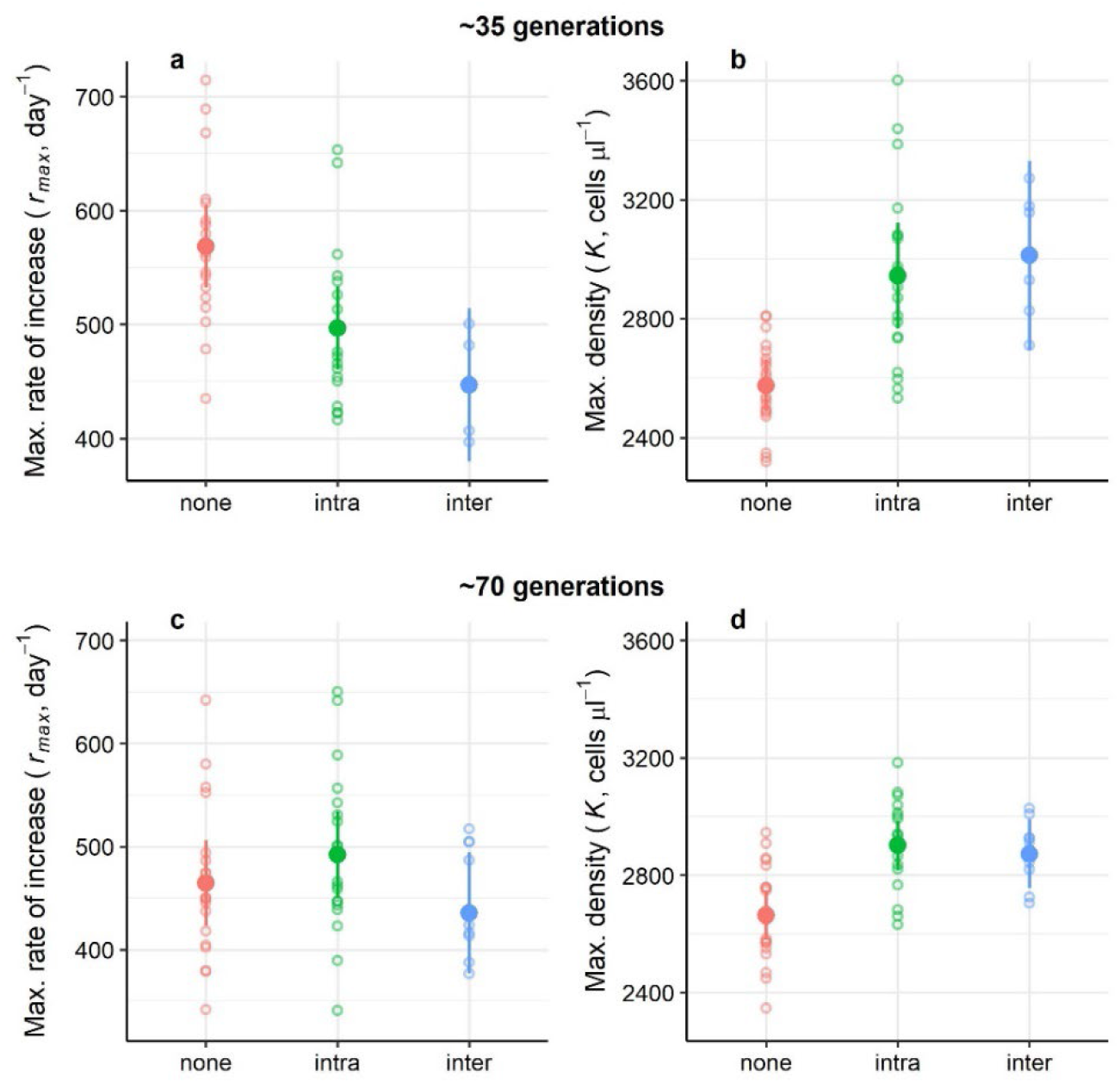
Populations that experienced competition had lower maximum population growth rates but greater carrying capacity in terms of cell numbers after 5 weeks of experimental evolution (~35 generations, top row). Differences in carrying capacity persisted after a further 35 generations but with no differences in growth rates between competition environments (bottom row). At either time, the type of competition experienced (intra- or inter-specific) did not affect demographic parameters.

We compared our quantitative predictions derived from metabolic theory with the observed scaling of each demographic parameter with cell size for the focal populations, and found remarkably strong congruence between the two for most parameters. We find that K_cells_ declines (c.f.: observed = –0.56; predicted = –0.60; F_1,48_ = 22.4, p < 0.001; CI: –0.79; –0.32; Fig. 6b) and K_bio_ increases with size as predicted by theory (c.f.: observed = 0.44; predicted = 0.40; F_1,48_ = 14.1, p < 0.001; Fig. 6c). But we find a slightly weaker than predicted relationship between *r*_max_ and size (c.f.: observed= 0.06; predicted = 0.13); this relationship is non-significant as the confidence intervals overlap zero (F_1,48_ = 0.02, p = 0.88; CI: –0.71; 0.83; Fig. 6a).

**Figure 6.**
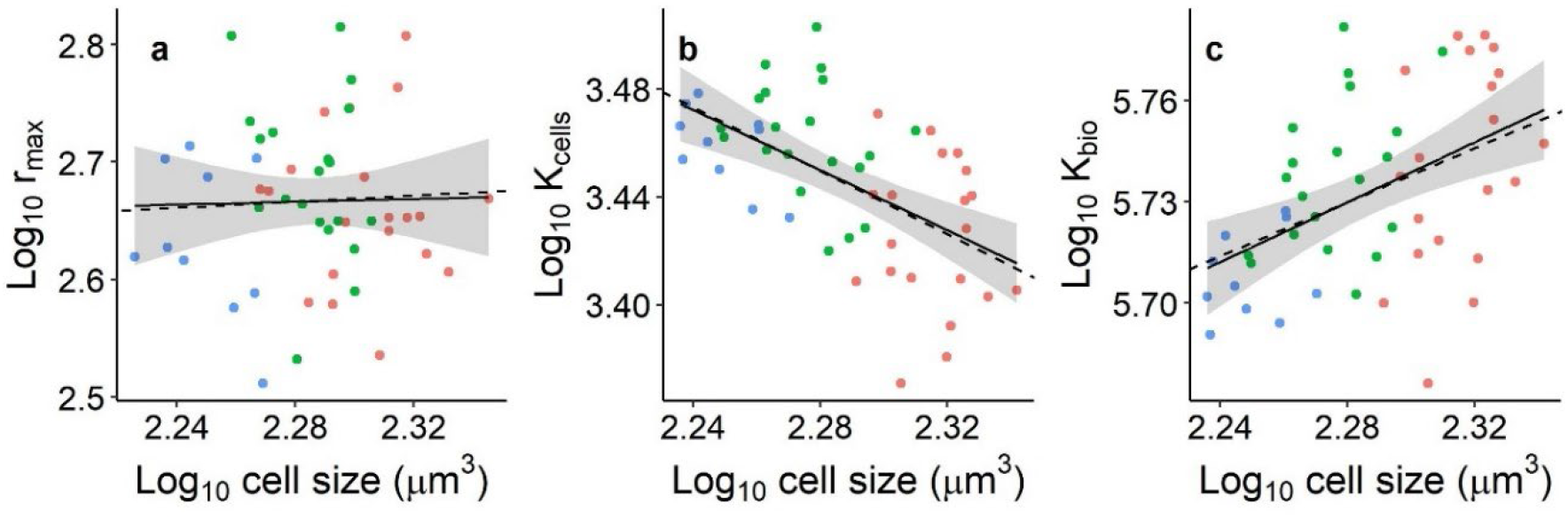
We find strong congruence between predictions from metabolic theory (broken lines) and demographic observations (solid lines). The scalings of max. growth rate (*r*_max_), max. population density (K_cells_) and max. biovolume (K_bio_) with cell size observed from our data after 70 generations are: *r*_max_ = observed 0.06 (CI: –0.71; 0.83); predicted 0.13; K_cells_ = observed –0.56 (CI: –0.79; –0.32); predicted –0.60; K_bio_ = observed 0.44 (CI: 0.21; 0.68); predicted 0.40. Scaling exponents are calculated for the 3 competition treatments together (blue = inter; green = intra; red = competition-free). We calculated the expected scaling of *r*_max_ as the average scaling observed over the first 5 days because population growth is determined early on (exponential growth phase); we calculated the expected scaling of K_cells_ and K_bio_ as the average scaling over the last 3 sampling days (12 to 16) when cultures were in stationary phase and because cell size stabilised from day 12 onwards (Fig. S2). We used the same range of days to calculate the average cell size for each replicate shown above (hence differences in size on x-axis between panel a and panels b-c).

## Discussion

We found that intra- and inter-specific competition drove the evolution of cell size and metabolism in the eukaryote *Dunaliella tertiolecta* in ways that metabolic theory anticipates. Because metabolism scaled hyper-allometrically with cell size in the focal lineages, the evolution of smaller sizes in response to competition reduced *per capita* resource demands (both in absolute and relative terms), such that cells achieved higher population densities but lower total biomass. These increases in population carrying capacity initially came at the expense of population growth rates – after only 35 generations of experimental evolution, the focal lineages experienced the classic trade-off between population growth rate and carrying capacity. However, through further evolution (~70 generations), competition-exposed lineages achieved Pareto improvements in demography: their max. population growth rates were comparable to those of competition-free lineages while also maintaining higher carrying capacities (Fig. 5). The evolution of greater metabolic flexibility appeared to enable this change; competition-exposed lineages were able to better downregulate their energy fluxes when resources were scare but strongly upregulated their metabolism when resources were abundant, thereby increasing both population growth rates and carrying capacity. This finding represents a novel departure from theory as the evolution of enhanced metabolic plasticity was unanticipated by any theory base but, in hindsight, seems almost inevitable. We eagerly await further studies that test the role of competition in driving metabolic plasticity in other systems. Enhancements in plasticity might explain why within-species patterns may not always reflect the same trade-offs observed among species ^14,26^.

Metabolism sets resource use and fuels growth, so metabolic rate should be under selection when competition is intense ^27^. Classic macroevolutionary patterns suggest that competition selects for efficiency which is often associated with larger body sizes and lower metabolic rates because metabolism scales hypo-allometrically in most metazoans ^28–30^. In contrast, our species exhibits hyper-allometric scaling of both photosynthesis (1.31) and respiration (1.28) with cell size so, while larger cells produce relatively more energy *per capita*, they are also both absolutely and relatively more expensive to maintain. Therefore, lineages exposed to competition evolved smaller, more resource-efficient cells. Interestingly, Long Term Experimental Evolution (LTEE) in *E. coli* yielded the opposite evolutionary trajectory of cell size to produce the same outcome – cells became larger over time to enhance resource-use efficiency ^25^. The trajectories of cell size in both studies actually accord with metabolic theory. In both, cells change size such that size-specific metabolic rates decrease (metabolism scales hypo-allometrically with size in the LTEE). Metabolic theory also predicts the relationships between size, density, and biomass in each study (based on species-specific metabolic scaling relationships). Based on our results and those of Marshall et al. ^25^, we would therefore predict that competition should select for smaller body size in species which show hyper-allometric metabolic scaling, but larger body size in species that show hypo-allometric scaling. For metazoans, most species show hypo-allometric metabolic scaling within species (the appropriate biological scale in this instance), whereas intraspecific metabolic scaling in unicellular organisms remains largely unexplored except for a few cases. We look forward to future studies that explore how metabolic scaling and competition interact in other species, particularly other microbial species given their pivotal role in driving biogeochemical cycles ^31,32^.

Our findings have some concerning implications regarding global change. Phytoplankton play a key role in the global carbon budget, accounting for ~50% of all carbon fixed annually; thus, any change in phytoplankton productivity will strongly influence global carbon fluxes ^33^. Increased water stratification due to ocean warming will intensify resource competition and decrease cell size within species ^34,35^. Our competition-exposed lineages evolved smaller cells in response to stronger resource competition – in that sense, our experimental evolution mimics the impacts of global change. We found that the net energy fluxes of the competition-exposed lineages were ~15% lower in the stationary phase than those of competition-free lineages (Fig. 2f). Hence the net carbon assimilation rate of our evolved lineages was systematically lower. If natural phytoplankton populations also evolve smaller cell sizes in response to warming and/or nutrient limitation (and initial studies suggest that they do) ^34^, then our results would imply that rates of carbon assimilation and phytoplankton biomass will decline with further global change, creating a positive feedback loop. While thermal adaptation might dampen the negative effects of warming on phytoplankton productivity ^36,37^, species interactions can increase the sensitivity of species to global change through resource competition ^7,38^ and alter their evolutionary rates in ways that are poorly resolved ^4,6^. Our results suggest that competition for resources might compound the impacts of global change by further selecting for smaller cell sizes and thriftier metabolisms that better tolerate competition but that may have undesirable impacts on carbon fixation.

Evolution happened rapidly in our system (within 70 generations). Such rapid evolution was likely fuelled by the very large population size of the founding populations (~5 million cells). These rates of evolution are comparable to those observed in other experimental evolution studies of phytoplankton where size and metabolic traits evolved within 100 generations ^37,39,40^, whereas it takes longer for evolution to manifest when populations are derived from a single clone ^41^. Overall, competition from a community exerted stronger effects on the evolution of these traits than intraspecific competition, possibly because diverse competitors consume available resources more rapidly and effectively ^42,43^. Thus, by posing a stronger selective pressure, competition from a community speeded up adaptation countering the negative effects of reduced population sizes due to stronger resource competition ^44^, similarly to what observed in bacteria ^4,5^. Still, we find remarkable consistency between the evolutionary responses to intra- and inter-specific competition, with both selecting for smaller, more efficient cells able to sustain greater maximum population densities. Carrying capacity (or equilibrium population size, K), rather than growth rate, is thus the quantity maximised by competition as predicted by life history theory for stable or low stochastic environments ^45–47^. Selection for r-K traits has been demonstrated in numerous systems within species ^27,46,48^ but few studies have demonstrated the same principle in response to interspecific competition. MacArthur & Wilson first hypothesized in 1967 that selection for K should also apply in a community when species compete for similar resources ^45^ but, to our best knowledge, these results are a first experimental demonstration that this demographic prediction of life history theory also holds under interspecific competition.

Since size and metabolism together determine resource access and use ^34,49,50^, changes in both traits might be a common feature of how species cope with shifts in competition and resource regimes. Both size and metabolic evolution might become more common, or rapid, as climate change, species introductions and biodiversity loss alter competition for resources ^51,52^. It seems likely that metabolic evolution will underpin many human-induced changes in resource regimes as well as mortality risks ^53^, and we look forward to future studies that search for signatures of shifts in metabolism associated with changes in competition regimes.

## Materials and Methods

### Overview of experimental set up

We tested the effects of competition on the metabolic fluxes, morphology, and demography of the eukaryotic unicellular alga *Dunaliella tertiolecta*, which we obtained from the CSIRO Australian National Algae Culture Collection (ANACC, strain CS-14). To manipulate the competitive environment, we enclosed the focal species in dialysis bags which were then placed in one of three environments for 10 weeks (1^st^ June to 10^th^ August 2021, ~70 generations): 1) surrounded by media with no competitors (competition-free), 2) by a population of conspecifics from the same strain (intraspecific competition) or 3) by a community of three other phytoplankton species (interspecific competition). For this latter treatment, the species were chosen to represent different sizes and phytoplankton groups: *Amphidinium carterae* (CS-740), *Phaeodactylum tricornutum* (CS-29) and *Nannochloropsis oculata* (CS-179) (Fig. 1, phase 1). Every week we batch-transferred both the focal species and the competitors (~7 generations). After 5 and 10 weeks (~35 and ~70 generations, respectively), we quantified changes in the traits of the focal species in common garden experiments (phase 2).

We sourced all phytoplankton species as 20 ml populations from CSIRO algal collection, Australia. We grew each population individually in 2 L glass bottles with standard f/2 enriched seawater medium enriched with silica for a month prior to the experiment, transferring each population into fresh media every week. The same medium was used throughout the experiment and common gardens. The medium was prepared with 0.45 μm filtered seawater and autoclaved following the recipe of Guillard & Ryther, 1962 ^54^. All experiments were performed in a temperature controlled room (20 ± 1 °C) on a 14:10 hours light:dark cycle and the cultures were grown under a light intensity of 115 ± 5 μmol photons m^−2^ s^−1^.

### Phase 1: evolution with competitors

The dialysis bags in which we enclosed the focal species enabled competition for light and nutrients but prevented cell mixing among phytoplankton species and exchange of bacteria (MWCO 12-14000 Da, pore size 24 Angstrom). Each bag contained a volume of 35 ml and was placed at the centre of a 500 ml glass jar assigned to one of the three competition treatments above (n = 20 for each competition treatment).

We started the experiment with an initial biovolume of 9.6 × 10^8^ μm^3^ of the focal species which corresponded 5 × 10^6^ cells. This large inoculum size meant that the founding populations likely contained a large amount of standing genetic variation. The glass jars containing the competitors were filled to 350 ml to completely submerge the dialysis bag. To maintain the same biovolume to media ratio, the initial biovolume of the competitors was 10 times that of the focal species (i.e. 9.6 × 10^9^ μm^3^). We added the three competitor species of the interspecific treatment in equal biovolumes. The jars of the control treatment (competition-free) were filled with media only.

We quantified biovolume as the product of cell density and cell size obtained for each species from two 10 μl samples, fixed with 1% Lugol’s solution. For each sample, we took 20 photos equally spaced around the cell-counting grid of a Neubauer counting chamber (ProSciTech, Australia) under an Olympus light microscope at 400× magnification. We analysed the images with ImageJ and Fiji software (version 2.0) ^55^ to quantify the number of cells (cells μl^−1^) and their size (μm, length and width; diameter for *Nannochloropsis*). We calculated cell volume by assigning to each species an approximate geometric shape ^56^ (prolate spheroid *V = (pi/6) × width^2^ × length* for all species except *Nannochloropsis* which we treated as a sphere *V = (pi/6) × diameter^3^*). Cell circularity is calculated by the Fiji software as *4 × pi × (area/perimeter^2^*) and ranges from 0 to 1 (perfect circle).

Each week we transferred the same initial biovolume of the focal species and the same initial biovolume of the competitors to a new, sterilized set of dialysis bag and jar. Cell densities, size and biovolume were determined as described above. We did not manipulate the relative abundance of species in the interspecific treatment. In weeks 3, 5 and 6 the cell densities of the focal species were very low in the interspecific treatment – had we reinoculated the same initial biovolume we would not have been able to add fresh medium. To avoid this nutrient limitation, we reinoculated only half of the initial biovolumes across all treatments. By week 6, we discarded 13 lineages of the focal species in the interspecific treatment because they were contaminated by one of the interspecific competitors (the diatom *Phaeodactylum tricornutum*). To continue the experiment, we established an additional lineage from each of the remaining seven replicates. By the end of the experiment, we lost 4 other replicates in the interspecific treatment because of *Phaeodactylum*’s contamination.

The batch-transfer approach meant that all populations in our experiment experienced fluctuating densities, but those surrounded by competitors always faced greater densities and thus competition. Shifts in densities are common in nature and mediate the balance between density-independent and density-dependent selection (r- and K-selection) ^45^. If stochasticity is low, life theory predicts that selection for K should prevail even under fluctuating conditions ^28,47,57^.

### Phase 2: common garden experiments

We quantified evolutionary changes in the metabolism, morphology and demography of the focal species in two common garden experiments; the first after five weeks of evolution with competitors (~35 generations) and the second after ten weeks (~70 generations). Before each common garden, we grew the focal species in a neutral environment for two generations to reduce any environmental conditioning (neutral selection). Both for the neutral selection and common garden, we inoculated an equal biovolume of each lineage in cell culture flasks (*n* = 20 for the no competition and intraspecific treatments, *n* = 6 and *n* = 10 for the interspecific treatment in the first and second common garden, respectively). Biovolume was determined as before from the average cell size and cell density of each lineage; we decided a priori to add 15 ml of the lineage with the limiting biovolume (~14 × 10^8^ μm^3^), back calculated the volumes of the other lineages and added 100 ml of media (f/2 + silica, as above) to all flasks. The two common gardens started on 8^th^ July and 12^th^ August 2021, respectively, and lasted 16 days, which was the approximate time needed for the cultures to reach carrying capacity – one of the key traits we wanted to phenotype. We discontinued the common garden after 16 days to avoid further evolution or adaptation by the focal lineages to the common garden conditions. Each day, we removed 10 ml from each culture flask for sampling and replaced it with fresh media. In the first common garden we sampled every day except on days 6, 10, 13 and 15; in the second we sampled every day except on days 7, 13 and 15.

### Trait measurements

Each sampling day, we fixed 1 ml sample with 1% Lugol’s solution to quantify the cell size (volume), shape (circularity) and abundance of each lineage as described above. We assessed changes in cell shape in addition to size because the shape of a cell can mediate access to resources and is thus an important component of fitness in unicellular organisms ^34,58–61^.

We measured oxygen evolution rates in 5 ml samples using 24-channel optical fluorescence oxygen readers (PreSens Sensor Dish Reader, SDR; AS-1 Scientific Wellington, New Zealand) following established protocols ^62,63^. Sensors were calibrated with 0% and 100% air saturation before the experiment. Net photosynthesis (oxygen production) was measured under the light intensity at which the cultures were grown for 20 minutes, followed by 1 hour in the dark to measure respiration rates. Thirteen blanks were filled with the media obtained from centrifuged samples (spun at 2,500 rpm for 10 min to separate the algae from the supernatant) to correct for background microbial activity since cultures were not axenic. Prior to measurements, samples were spiked with 50 μl of sodium bicarbonate stock for a final concentration of 2 mM sodium bicarbonate to avoid carbon limitation.

The change in percentage oxygen saturation was calculated with linear regressions using the LoLinR package ^64^. The rate of photosynthesis or respiration of the whole sample (VO_2_; units μmol O_2_/min) was then measured as VO_2_ = 1 × ((m_a_ – m_b_)/100 × VβO_2_) following ^65^, where m_a_ is the rate of change of O_2_ saturation in each sample (min^−1^), mb is the mean O_2_ saturation across all blanks (min^−1^), V is the sample volume (0.005 L) and VβO_2_ is the oxygen capacity of air-saturated seawater at 20°C and 35 ppt salinity (225 μmol O_2_/L). The first three minutes of measurements in the light were discarded for all samples. Respiration rates were calculated after 15 minutes of dark when oxygen levels started to show a linear decline. Photosynthesis and respiration (μmol O_2_/min) were converted to calorific energy (J/min) using the conversion factor of 0.512 J/μmol O_2_ to estimate energy production and energy consumption, respectively ^66^.

### Statistical analyses

All analyses and plots were done in RStudio (version 4.1.3), separately for the two common gardens, using the main packages nlme ^67^, lme4 ^68^, car ^69^, and ggplot2 ^70^.

#### Morphology

We assessed differences in cell morphology (size and shape) with linear mixed models including competition treatment (3 levels) and time (day) as categorical predictors, and lineage identity as random intercept to account for repeated measures; time was considered categorical because the relationship with cell morphology was non-linear. We take day 3, during the exponential growth phase, as a reference to report post hoc results on differences in cell size and shape in the main text, and we report post hoc results for each day in supplements.

#### Metabolism

While oxygen rates increased linearly with biovolume over the first few days, this relationship broke down as biovolume increased (over ~3 μm^3^/μl) (Fig. S4). Log-transformation did not result in linearity across the entire range of biovolume. Therefore, we analysed oxygen rates separately for the first part of common gardens during the exponential growth phase (from day 1 to day 6 included), and for the second part during the stationary phase (day 7 to 16). We used linear mixed models with biovolume (covariate) and competition treatment as predictors, and lineage identity as random intercept for the data of the second common garden to account for repeated measures. Because variances were heterogenous for the photosynthesis data during the growth phase we used a generalised linear mixed model including competition-specific variances. Finally, we integrated photosynthesis and respiration rates over a 24-hour period to estimate net energy production of the whole sample (J/day), as 14 hr of energy produced through net photosynthesis minus 10 hr of respiration using rates predicted from the mixed models above.

We could not use mixed models on the oxygen evolution data from the first common garden because of a singular fit error likely due to the lower replication of the interspecific competition treatment (*n* = 6). Therefore, we used linear models with biovolume (covariate) and competition treatment as predictors (i.e. without including lineage identity). While this is not ideal, we repeated the analyses for the second common garden with the same, simple linear models and obtained the same results of mixed models (results not reported here), suggesting that oxygen rates are not strongly affected by lineage identity. Nonetheless, we report the results on oxygen rates for the first common garden as a supplementary figure. For all analyses of oxygen rates, data were not transformed but biovolume was rescaled to 10^−5^ μm^3^ μl^−1^. Interactions between biovolume and competition were removed when p > 0.25 or if the interaction was not significant and the model with interaction did not perform better than the simpler model compared by AIC and anova.

#### Demography

To test differences in the maximum rates of increase (*r*_max_) and maximum values (K_cells_) of cell density we followed a three-step approach (described in detail in Ghedini et al. ^63^, adapted from Malerba et al. ^71^). First, we fitted four growth models to each individual replicate lineage and chose the best-fitting model among the four candidates to describe changes in the cell density (cells μl^−1^) of each culture over time. We used AIC to determine which growth model best described the dynamics of a culture and successful convergence was ensured for all best-fitting models. The four models were: a logistic-type sinusoidal growth model with lower asymptote forced to 0 (i.e. three-parameter logistic curve), a logistic-type sinusoidal growth model with non-zero lower asymptote (i.e. four-parameter logistic curve), a Gompertz-type sinusoidal growth model (i.e. three-parameter Gompertz curve) and a modified Gompertz-type sinusoidal growth model including population decline after reaching a maximum (i.e. four-parameter Gompertz-like curve including mortality). Second, we used the best-fitting model to estimate growth parameters (i.e. *r*_max_ and K_cells_) for each culture. From each nonlinear curve, we extracted the maximum predicted value of population density (K_cells_, cells μl^−1^). From the first derivative of the curve, we extracted the maximum rate of population increase (*r*_max_, unit: day^−1^). Third, we used an analysis of covariance to evaluate the influence of competition on each parameter, using a linear model including the initial cell density estimated from the previous step as a covariate and competition environment as a factor (three levels). The estimates of K_cells_ of the first common garden had heterogeneous variances among treatments, so we used generalized least squares models (instead of linear models) including treatment-specific variance for each level of competition treatment (varIdent function in R). We then estimated and plotted least square means and 95% confidence intervals using Tukey p-value adjustment for comparing three estimates.

#### Scaling of demographic parameters with cell size

We determined the scaling of metabolism with cell size for our evolved populations after ten weeks of evolution (~70 generation). For this assessment we combined the data across the three competition treatments to assess a wider range of size. We calculated the scaling of respiration and photosynthesis, separately, from linear models including cell size and experiment day as numerical predictors. Oxygen rates and cell size were log10-transformed. Results did not qualitatively change when we analysed each competition treatment separately, as each of them scaled > 1 and declined over time (Table S4).

Based on these results (effects of size and experiment day), we then calculated the expected scaling of respiration with size for any given day. According to metabolic theory, the cost of production is directly proportional to cell size (scales with size at 1) ^17,25^, so demographic parameters should scale with size at:

1. *r*_max_ = M^B^_exponential_/M^1^ = M^B-1^
2. K_cells_ = M^0^/M^B^_stationary_ = M^−B^
3. K_bio_ = M × K_cells_ = M × M^−B^ = M^1-B^

We tested these predictions by comparing the predicted metabolic scaling values for the growth phase and stationary phase to the observed metabolic scalings from our data. We used early (days 1-5) respiratory scaling values for expectations about *r*_max_ because it is mostly determined by growth rates early on (B_exponential_ = 1.13). We used late (days 12-16) respiratory scaling values for expectations about K_cells_ and K_bio_ (B_stationary_ = 0.60) because cultures reached carrying capacity in the latter part of the common garden and because cell size (which affects estimates of K_bio_) stabilised around day 12 (Fig. S2). We estimated maximum total biomass (K_bio_) as the product of K_cells_ obtained from the population growth models and the average cell size for each replicate over the last three sampling days (day 12 to 16), when cell size stabilised. Finally, we determined the empirical scaling of each demographic parameter with cell size across the focal populations, using the average cell size calculated over the same intervals above (i.e., over the first 5 days for *r*_max_, between day 12 and 16 for K_cells_ and K_bio_).

## Acknowledgements

We thank Mike McDonald and John DeLong for valuable comments on earlier versions of the manuscript. We also thank Jiaye Qin for assistance in data collection and the Quantitative Biology Unit at the Instituto Gulbenkian de Ciência for statistical advice. This work was supported by the Australian Research Council through a DECRA fellowship (DE190100660) and by a fellowship (LCF/BQ/PI21/11830001) from “la Caixa” Foundation (ID 100010434) and the European Union’s Horizon 2020 research and innovation programme under the Marie Skłodowska Curie grant agreement No 847648 to GG.

