## Supplementary Information for "Metabolic evolution in response to interspecific competition in a eukaryote"

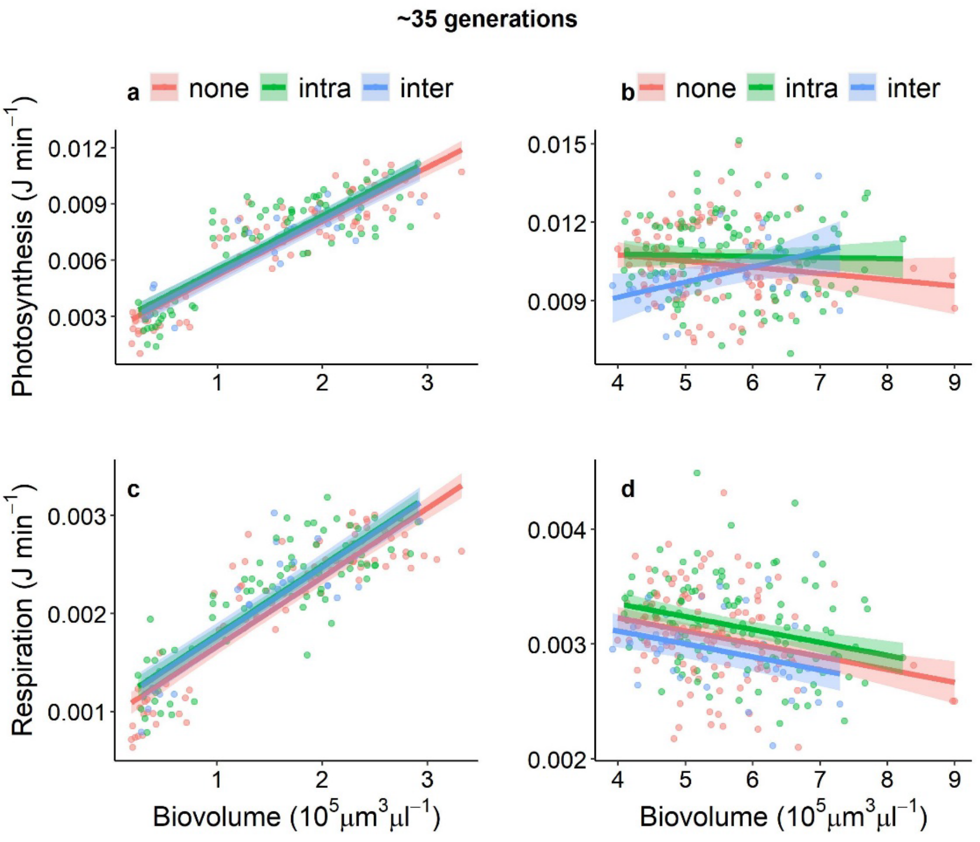

**Figure S1.** Population rates of photosynthesis and respiration as a function of population biovolume during the exponential growth phase (left) and stationary phase (right) with no competitors ("none"), intraspecific ("intra") or interspecific competitors ("inter"). The focal species shows signs of metabolic evolution after ~35 generations (5 weeks) with competitors but only when metabolism is measured under resource limitation (i.e. stationary phase).

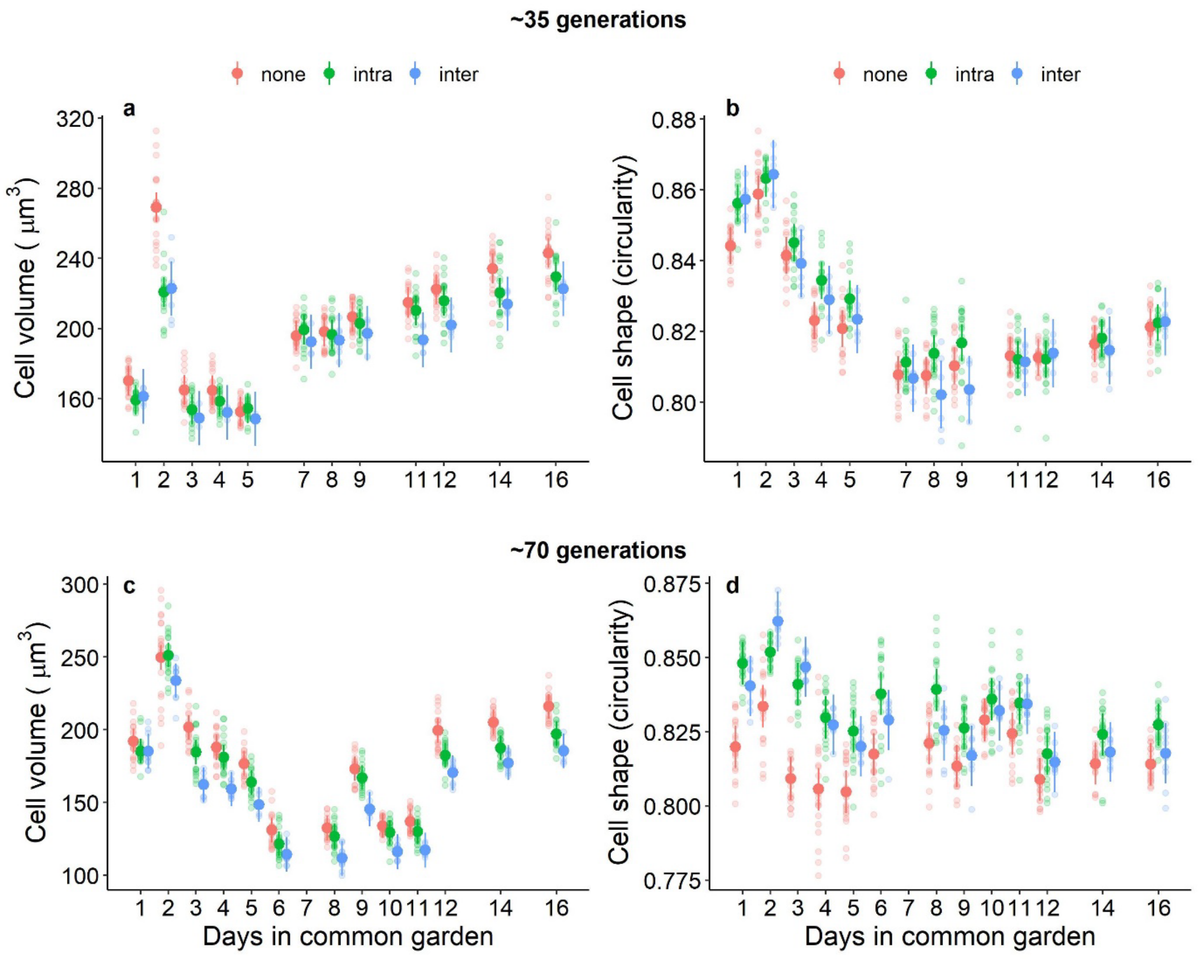

**Figure S2.** Cell size (cell volume, left) and shape (circularity, right) of the focal species evolve in response to competition, and these evolutionary responses strengthen with longer-term evolution with competitors (35 generations, top; 70 generations, bottom). Populations exposed to competitors evolved smaller cell sizes, and the reduction in size was stronger in response to inter-specific competitors (a, c). Changes in cell size are accompanied by changes in shape – overall, the smaller cells exposed to competitors tend to be rounder. See Table S3 for post hoc comparisons for each day.

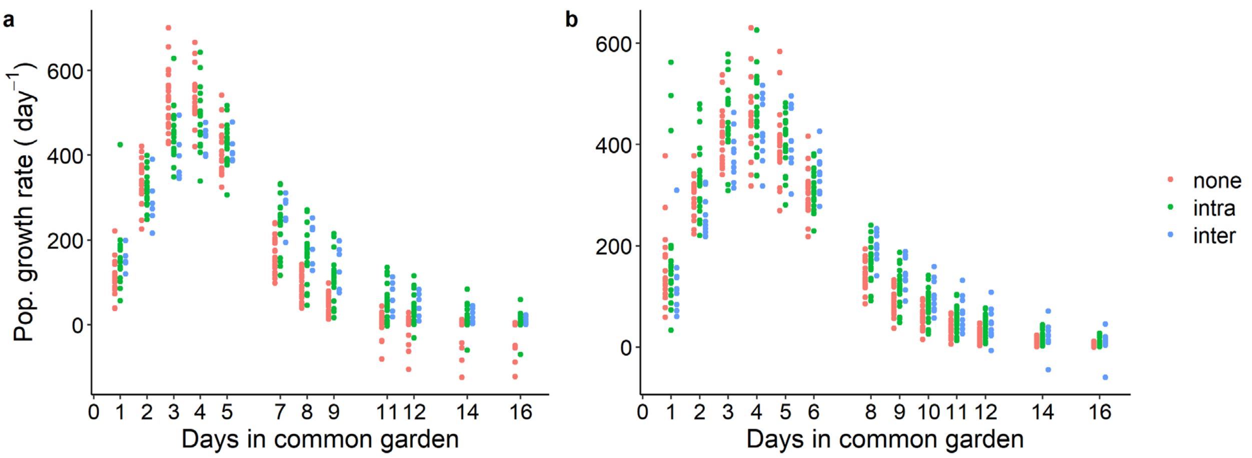

**Fig. S3.** Population growth rate of the focal lineages for each day of common garden experiments after ~35 (a) and ~70 generations (b) of experimental evolution. In both cases, populations that evolved with intra- or inter-specific competition sustained greater population growth as they approached carrying capacity than populations evolved without competitors.

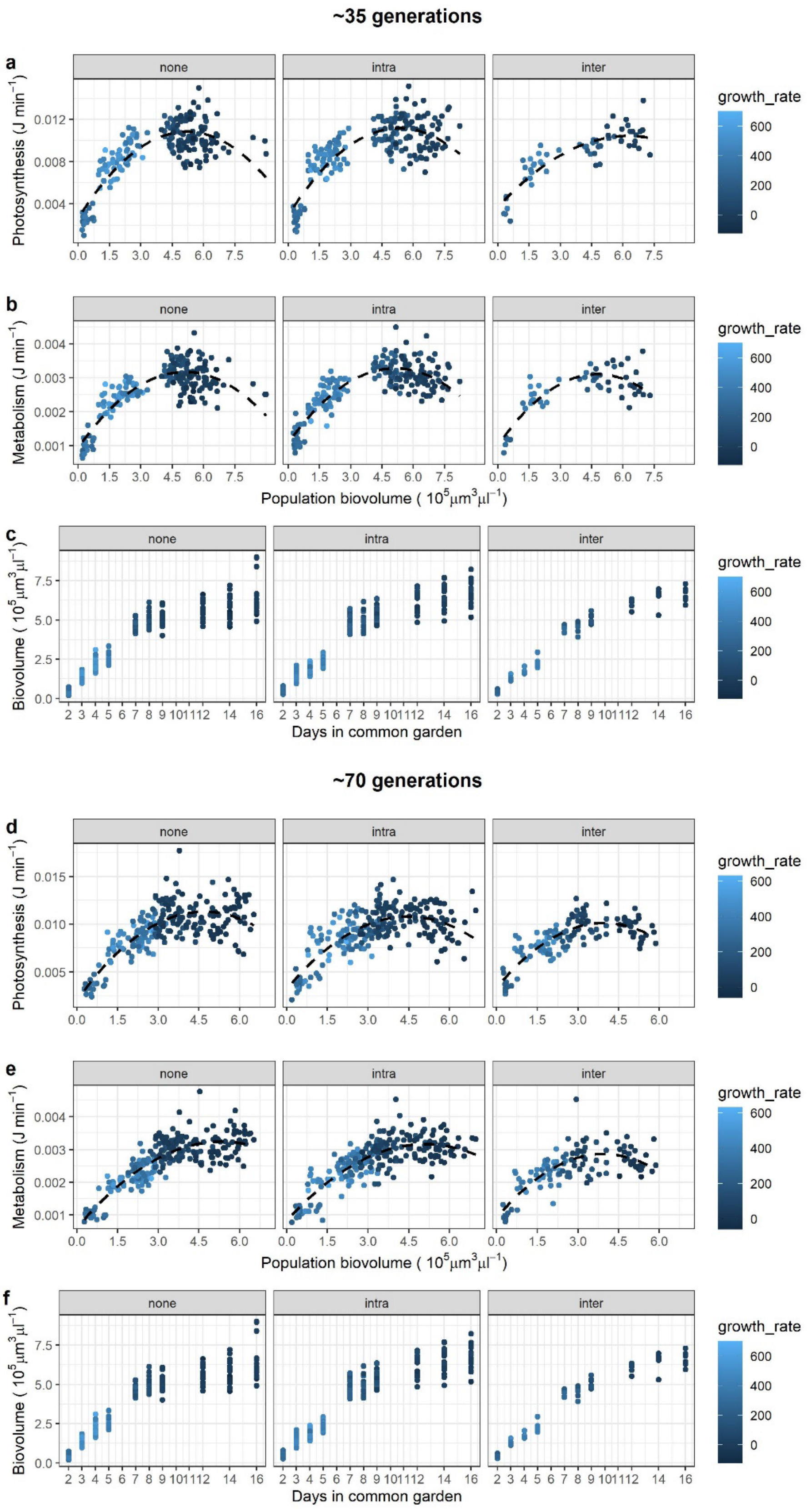

**Fig. S4.** Photosynthesis and respiration increased nearly linearly with biovolume in the first part of common gardens (exponential growth phase) but this relationship broke down as populations approached carrying capacity and growth slowed. This transition occurred at approximately 3 × $10^5$ $\mu m^3$ $\mu l^{-1}$ of biovolume, which most cultures reached around day 7.

**Table S1.** Linear models testing the effects of competition and biovolume (covariate, $10^{-5}$ µm$^3$/µl) on photosynthesis and respiration during the exponential growth phase or stationary phase after ~35 and ~70 generations of experimental evolution. We used simple linear models at 35 generations and mixed models at 70 generations (see Materials and Methods); the latter included lineage identity as a random intercept, and competition-specific variances for photosynthesis data during the exponential growth phase because variances were heterogenous. P values < 0.05 are in bold.

| **~35 generations** | | | | | |
|---|---|---|---|---|---|
| *Exponential phase* | | | | | |
| **Photosynthesis** | Df | Sum sq | Mean sq | F value | Pr(>F) |
| Biovolume | 1 | 0.00099 | 0.000994 | 608.97 | **<0.0001** |
| Competition | 2 | 0.0000036 | 0.000002 | 1.11 | 0.33 |
| Residuals | 178 | 0.0002908 | 0.000002 | | |
| **Respiration** | Df | Sum sq | Mean sq | F value | Pr(>F) |
| Biovolume | 1 | $5.9 \times 10^{-5}$ | $5.9 \times 10^{-5}$ | 542.61 | **<0.0001** |
| Competition | 2 | $6.0 \times 10^{-7}$ | $3.0 \times 10^{-7}$ | 2.75 | 0.067 |
| Residuals | 176 | $1.9 \times 10^{-5}$ | $1.1 \times 10^{-7}$ | | |
| *Stationary phase* | | | | | |
| **Photosynthesis** | Df | Sum sq | Mean sq | F value | Pr(>F) |
| Biovolume | 1 | 0.00000002 | $1.5 \times 10^{-8}$ | 0.007 | 0.93 |
| Competition | 2 | 0.000014 | $7.1 \times 10^{-6}$ | 3.372 | **0.04** |
| Biovolume × Competition | 2 | 0.00002 | $8.1 \times 10^{-6}$ | 3.847 | **0.02** |
| Residuals | 270 | 0.00057 | $2.1 \times 10^{-6}$ | | |
| contrast | estimate | SE | df | t-ratio | p-value |
| none - intra | -0.000193 | 0.000203 | 270 | -0.95 | 0.61 |
| none - inter | -0.000815 | 0.000294 | 270 | -2.77 | **0.02** |
| intra - inter | -0.000623 | 0.000289 | 270 | -2.16 | 0.08 |
| **Respiration** | Df | Sum Sq | Mean Sq | F value | Pr(>F) |
| Biovolume | 1 | $2.5 \times 10^{-6}$ | $2.5 \times 10^{-6}$ | 17.998 | **<0.0001** |
| Competition | 2 | $1.8 \times 10^{-6}$ | $9.2 \times 10^{-7}$ | 6.563 | **0.002** |
| Residuals | 272 | $3.8 \times 10^{-5}$ | $1.4 \times 10^{-7}$ | | |
| contrast | estimate | SE | df | t-ratio | p-value |
| none - intra | -0.00012 | $4.9 \times 10^{-5}$ | 272 | -2.51 | **0.03** |
| none - inter | 0.00011 | $7.1 \times 10^{-5}$ | 272 | 1.60 | 0.25 |
| intra - inter | 0.00024 | $7.1 \times 10^{-5}$ | 272 | 3.32 | **0.01** |

*Continues on next page*

| ~70 generations | | | | | | |
|---|---|---|---|---|---|---|
| | | | *Exponential phase* | | | |
| **Photosynthesis** | Sum sq | Mean sq | NumDF | DenDF | F value | Pr(>F) |
| Biovolume | 0.00058 | 0.00058 | 1 | 207.64 | 296.76 | **<0.0001** |
| Competition | $1.9 \times 10^{-5}$ | $9.5 \times 10^{-6}$ | 2 | 241.46 | 4.80 | **0.009** |
| Biovol. × Competition | $1.4 \times 10^{-5}$ | $7.1 \times 10^{-6}$ | 2 | 209.55 | 3.59 | **0.03** |
| *Slope estimates* | estimate | SE | df | lower | upper CL | |
| none | 0.0025 | 0.00017 | 212 | 0.0022 | 0.0029 | |
| intra | 0.0019 | 0.00017 | 219 | 0.0016 | 0.00225 | |
| inter | 0.0020 | 0.00029 | 204 | 0.0014 | 0.00256 | |
| *Contrast* | estimate | SE | df | t-ratio | p-value | |
| none - intra | $6.2 \times 10^{-4}$ | 0.00024 | 215 | 2.57 | **0.03** | |
| none - inter | $5.5 \times 10^{-4}$ | 0.00034 | 206 | 1.64 | 0.23 | |
| intra - inter | $-6.4 \times 10^{-5}$ | 0.00033 | 207 | -0.19 | 0.98 | |
| **Respiration** | Sum sq | Mean sq | NumDF | DenDF | F value | Pr(>F) |
| Biovolume | $5.7 \times 10^{-5}$ | $5.7 \times 10^{-5}$ | 1 | 216.34 | 502.13 | **<0.0001** |
| Competition | $2.5 \times 10^{-6}$ | $1.2 \times 10^{-6}$ | 2 | 46.14 | 10.86 | **0.0001** |
| *Contrast* | estimate | SE | df | t-ratio | p-value | |
| none - intra | -0.0001 | $4.9 \times 10^{-5}$ | 46.9 | -2.69 | **0.03** | |
| none - inter | -0.0003 | $6.1 \times 10^{-5}$ | 50.4 | -4.56 | **0.0001** | |
| intra - inter | -0.0002 | $6.1 \times 10^{-5}$ | 50.5 | -2.41 | **0.0507** | |

| *Stationary phase* | | | | |
|---|---|---|---|---|
| **Photosynthesis** | numDF | denDF | F-value | p-value |
| Intercept | 1 | 299 | 6808.12 | **<0.0001** |
| Biovolume | 1 | 299 | 4.74 | **0.0302** |
| Competition | 2 | 47 | 6.75 | **0.0027** |

| *Contrast* | estimate | SE | df | t-ratio | p-value |
|---|---|---|---|---|---|
| none - intra | 0.00042 | 0.00029 | 47 | 1.46 | 0.32 |
| none - inter | 0.00125 | 0.00034 | 47 | 3.67 | **0.002** |
| intra - inter | 0.00083 | 0.00034 | 47 | 2.43 | **0.049** |

| **Respiration** | Sum sq | Mean sq | NumDF | DenDF | F value | Pr(>F) |
|---|---|---|---|---|---|---|
| Biovolume | $9.6 \times 10^{-7}$ | $9.6 \times 10^{-7}$ | 1 | 320 | 6.46 | **0.01** |
| Competition | $4.7 \times 10^{-6}$ | $2.3 \times 10^{-6}$ | 2 | 47 | 15.83 | **<0.0001** |

| *Contrast* | estimate | SE | df | t-ratio | p-value |
|---|---|---|---|---|---|
| none - intra | $6.2 \times 10^{-5}$ | $5.2 \times 10^{-5}$ | 46.9 | 1.174 | 0.47 |
| none - inter | $3.6 \times 10^{-4}$ | $6.9 \times 10^{-5}$ | 48.7 | 5.507 | **<0.0001** |
| intra - inter | $2.95 \times 10^{-4}$ | $6.5 \times 10^{-5}$ | 48.7 | 4.559 | **0.0001** |

**Table S2.** Linear mixed models testing the effects of competition and time (days in common garden) on cell size and shape after ~35 and 70 generations of evolution with competitors. The models included replicate ID as a random intercept and time as a categorical variable because the relationship with cell size/shape was not linear. t-tests use Satterthwaite's method. P values < 0.05 are in bold.

| ~35 generations | | | | | | |
|---|---|---|---|---|---|---|
| **Cell size** | Sum Sq | Mean Sq | Num df | Den df | F value | p-value |
| Competition | 4982 | 2491 | 2 | 43 | 24.64 | **< 0.0001** |
| Time | 343554 | 31232 | 11 | 473 | 308.98 | **< 0.0001** |
| Competition × Time | 22134 | 1006 | 22 | 473 | 9.95 | **< 0.0001** |
| **Cell shape** | | | | | | |
| Competition | 0.00089 | 0.0005 | 2 | 43 | 11.27 | **0.0001** |
| Time | 0.12225 | 0.0111 | 11 | 473 | 278.52 | **< 0.0001** |
| Competition × Time | 0.00329 | 0.0002 | 22 | 473 | 3.75 | **< 0.0001** |
| **~70 generations** | | | | | | |
| **Cell size** | Sum Sq | Mean Sq | Num df | Den df | F value | p-value |
| Competition | 15734 | 7867 | 2 | 47 | 76.07 | **< 0.0001** |
| Time | 708876 | 59073 | 12 | 564 | 571.24 | **< 0.0001** |
| Competition × Time | 9069 | 378 | 24 | 564 | 3.65 | **< 0.0001** |
| **Cell shape** | | | | | | |
| Competition | 0.008824 | 0.0044 | 2 | 47 | 65.24 | **< 0.0001** |
| Time | 0.051959 | 0.0043 | 12 | 564 | 64.03 | **< 0.0001** |
| Competition × Time | 0.013793 | 0.0006 | 24 | 564 | 8.50 | **< 0.0001** |

**Table S3.** Post hoc comparisons (estimated marginal means) between competition treatments for cell size and shape for each day of common gardens. Time (exp day) is treated as a categorical variable. None = no competition. Intra = intraspecific competition. Inter = interspecific competition. P values < 0.05 are in bold.

| **Cell size after ~35 generations** | | | | | |
|---|---|---|---|---|---|
| contrast | estimate | SE | df | t-ratio | p-value |
| exp_day = 1: | | | | | |
| none - intra | 10.92 | 3.52 | 377 | 3.104 | **0.006** |
| none - inter | 8.81 | 5.18 | 377 | 1.702 | 0.206 |
| intra - inter | -2.11 | 5.18 | 377 | -0.407 | 0.91 |
| exp_day = 2: | | | | | |
| none - intra | 48.4 | 3.52 | 377 | 13.761 | **<.0001** |
| none - inter | 46.39 | 5.18 | 377 | 8.959 | **<.0001** |
| intra - inter | -2.02 | 5.18 | 377 | -0.389 | 0.92 |
| exp_day = 3: | | | | | |
| none - intra | 11.15 | 3.52 | 377 | 3.169 | **0.005** |
| none - inter | 16.14 | 5.18 | 377 | 3.118 | **0.006** |
| intra - inter | 5 | 5.18 | 377 | 0.965 | 0.60 |
| exp_day = 4: | | | | | |
| none - intra | 6.15 | 3.52 | 377 | 1.747 | 0.19 |
| none - inter | 12.56 | 5.18 | 377 | 2.425 | **0.042** |
| intra - inter | 6.41 | 5.18 | 377 | 1.239 | 0.43 |
| exp_day = 5: | | | | | |
| none - intra | -1.84 | 3.52 | 377 | -0.524 | 0.86 |
| none - inter | 4.1 | 5.18 | 377 | 0.792 | 0.71 |
| intra - inter | 5.94 | 5.18 | 377 | 1.148 | 0.49 |
| exp_day = 7: | | | | | |
| none - intra | -3.5 | 3.52 | 377 | -0.996 | 0.58 |
| none - inter | 3.29 | 5.18 | 377 | 0.636 | 0.80 |
| intra - inter | 6.8 | 5.18 | 377 | 1.313 | 0.39 |
| exp_day = 8: | | | | | |
| none - intra | 1.59 | 3.52 | 377 | 0.453 | 0.89 |
| none - inter | 4.7 | 5.18 | 377 | 0.907 | 0.64 |
| intra - inter | 3.1 | 5.18 | 377 | 0.6 | 0.82 |
| exp_day = 9: | | | | | |
| none - intra | 3.79 | 3.52 | 377 | 1.078 | 0.53 |
| none - inter | 9.48 | 5.18 | 377 | 1.831 | 0.16 |
| intra - inter | 5.69 | 5.18 | 377 | 1.098 | 0.52 |
| exp_day = 11: | | | | | |

| | | | | | |
|---|---|---|---|---|---|
| none - intra | 4.66 | 3.52 | 377 | 1.323 | 0.38 |
| none - inter | 21.41 | 5.18 | 377 | 4.135 | **0.0001** |
| intra - inter | 16.76 | 5.18 | 377 | 3.236 | **0.0038** |
| exp_day = 12: | | | | | |
| none - intra | 6.54 | 3.52 | 377 | 1.858 | 0.15 |
| none - inter | 20.27 | 5.18 | 377 | 3.915 | **0.0003** |
| intra - inter | 13.73 | 5.18 | 377 | 2.653 | **0.0226** |
| exp_day = 14: | | | | | |
| none - intra | 13.7 | 3.52 | 377 | 3.895 | **0.0003** |
| none - inter | 20.05 | 5.18 | 377 | 3.872 | **0.0004** |
| intra - inter | 6.35 | 5.18 | 377 | 1.226 | 0.44 |
| exp_day = 16: | | | | | |
| none - intra | 13.5 | 3.52 | 377 | 3.838 | **0.0004** |
| none - inter | 20.31 | 5.18 | 377 | 3.923 | **0.0003** |
| intra - inter | 6.81 | 5.18 | 377 | 1.316 | 0.39 |
| **Cell shape after ~35 generations** | | | | | |
| contrast | estimate | SE | df | t-ratio | p-value |
| exp_day = 1: | | | | | |
| none - intra | -0.012 | 0.00217 | 410 | -5.528 | **<.0001** |
| none - inter | -0.01314 | 0.0032 | 410 | -4.113 | **0.0001** |
| intra - inter | -0.00114 | 0.0032 | 410 | -0.357 | 0.93 |
| exp_day = 2: | | | | | |
| none - intra | -0.00443 | 0.00217 | 410 | -2.041 | 0.10 |
| none - inter | -0.00556 | 0.0032 | 410 | -1.74 | 0.19 |
| intra - inter | -0.00113 | 0.0032 | 410 | -0.353 | 0.93 |
| exp_day = 3: | | | | | |
| none - intra | -0.00356 | 0.00217 | 410 | -1.637 | 0.23 |
| none - inter | 0.002248 | 0.0032 | 410 | 0.703 | 0.76 |
| intra - inter | 0.005802 | 0.0032 | 410 | 1.816 | 0.16 |
| exp_day = 4: | | | | | |
| none - intra | -0.01129 | 0.00217 | 410 | -5.198 | **<0.0001** |
| none - inter | -0.00585 | 0.0032 | 410 | -1.83 | 0.16 |
| intra - inter | 0.005437 | 0.0032 | 410 | 1.702 | 0.21 |
| exp_day = 5: | | | | | |
| none - intra | -0.00833 | 0.00217 | 410 | -3.837 | **0.0004** |
| none - inter | -0.00264 | 0.0032 | 410 | -0.825 | 0.69 |
| intra - inter | 0.005693 | 0.0032 | 410 | 1.782 | 0.18 |
| exp_day = 7: | | | | | |

| | | | | | |
|---|---|---|---|---|---|
| none - intra | -0.00359 | 0.00217 | 410 | -1.653 | 0.22 |
| none - inter | 0.000991 | 0.0032 | 410 | 0.31 | 0.95 |
| intra - inter | 0.00458 | 0.0032 | 410 | 1.433 | 0.32 |
| exp_day = 8: | | | | | |
| none - intra | -0.0062 | 0.00217 | 410 | -2.856 | **0.013** |
| none - inter | 0.005395 | 0.0032 | 410 | 1.688 | 0.21 |
| intra - inter | 0.011595 | 0.0032 | 410 | 3.629 | **0.0009** |
| exp_day = 9: | | | | | |
| none - intra | -0.00651 | 0.00217 | 410 | -3 | **0.008** |
| none - inter | 0.006723 | 0.0032 | 410 | 2.104 | 0.09 |
| intra - inter | 0.013236 | 0.0032 | 410 | 4.142 | **0.0001** |
| exp_day = 11: | | | | | |
| none - intra | 0.001112 | 0.00217 | 410 | 0.512 | 0.87 |
| none - inter | 0.001812 | 0.0032 | 410 | 0.567 | 0.84 |
| intra - inter | 0.0007 | 0.0032 | 410 | 0.219 | 0.97 |
| exp_day = 12: | | | | | |
| none - intra | 0.000446 | 0.00217 | 410 | 0.206 | 0.98 |
| none - inter | -0.00125 | 0.0032 | 410 | -0.393 | 0.92 |
| intra - inter | -0.0017 | 0.0032 | 410 | -0.532 | 0.86 |
| exp_day = 14: | | | | | |
| none - intra | -0.00162 | 0.00217 | 410 | -0.746 | 0.74 |
| none - inter | 0.001863 | 0.0032 | 410 | 0.583 | 0.83 |
| intra - inter | 0.003484 | 0.0032 | 410 | 1.09 | 0.52 |
| exp_day = 16: | | | | | |
| none - intra | -0.00114 | 0.00217 | 410 | -0.526 | 0.86 |
| none - inter | -0.0015 | 0.0032 | 410 | -0.468 | 0.89 |
| intra - inter | -0.00035 | 0.0032 | 410 | -0.111 | 0.99 |
| **Cell size after ~70 generations** | | | | | |
| contrast | estimate | SE | df | t-ratio | p-value |
| exp_day = 1: | | | | | |
| none - intra | 6.9077 | 3.47 | 492 | 1.989 | 0.12 |
| none - inter | 6.8891 | 4.25 | 492 | 1.62 | 0.24 |
| intra - inter | -0.0186 | 4.25 | 492 | -0.004 | 1 |
| exp_day = 2: | | | | | |
| none - intra | -1.5014 | 3.47 | 492 | -0.432 | 0.90 |
| none - inter | 15.8582 | 4.25 | 492 | 3.729 | **0.0006** |
| intra - inter | 17.3596 | 4.25 | 492 | 4.082 | **0.0002** |
| exp_day = 3: | | | | | |

| | | | | | |
|---|---|---|---|---|---|
| none - intra | 17.124 | 3.47 | 492 | 4.932 | **<0.0001** |
| none - inter | 39.5429 | 4.25 | 492 | 9.299 | **<0.0001** |
| intra - inter | 22.4189 | 4.25 | 492 | 5.272 | **<0.0001** |
| exp_day = 4: | | | | | |
| none - intra | 6.8958 | 3.47 | 492 | 1.986 | 0.12 |
| none - inter | 28.7018 | 4.25 | 492 | 6.749 | **<0.0001** |
| intra - inter | 21.8059 | 4.25 | 492 | 5.128 | **<0.0001** |
| exp_day = 5: | | | | | |
| none - intra | 12.8299 | 3.47 | 492 | 3.695 | **0.0007** |
| none - inter | 28.0425 | 4.25 | 492 | 6.594 | **<0.0001** |
| intra - inter | 15.2126 | 4.25 | 492 | 3.577 | **0.0011** |
| exp_day = 6: | | | | | |
| none - intra | 9.5038 | 3.47 | 492 | 2.737 | **0.02** |
| none - inter | 16.7982 | 4.25 | 492 | 3.95 | **0.0003** |
| intra - inter | 7.2944 | 4.25 | 492 | 1.715 | 0.20 |
| exp_day = 8: | | | | | |
| none - intra | 5.7617 | 3.47 | 492 | 1.659 | 0.22 |
| none - inter | 20.7359 | 4.25 | 492 | 4.876 | **<0.0001** |
| intra - inter | 14.9742 | 4.25 | 492 | 3.521 | **0.0014** |
| exp_day = 9: | | | | | |
| none - intra | 6.0451 | 3.47 | 492 | 1.741 | 0.19 |
| none - inter | 27.6995 | 4.25 | 492 | 6.514 | **<0.0001** |
| intra - inter | 21.6545 | 4.25 | 492 | 5.092 | **<0.0001** |
| exp_day = 10: | | | | | |
| none - intra | 4.592 | 3.47 | 492 | 1.323 | 0.38 |
| none - inter | 17.8785 | 4.25 | 492 | 4.204 | **0.0001** |
| intra - inter | 13.2865 | 4.25 | 492 | 3.124 | **0.0054** |
| exp_day = 11: | | | | | |
| none - intra | 6.9309 | 3.47 | 492 | 1.996 | 0.11 |
| none - inter | 19.679 | 4.25 | 492 | 4.628 | **<0.0001** |
| intra - inter | 12.7481 | 4.25 | 492 | 2.998 | **0.008** |
| exp_day = 12: | | | | | |
| none - intra | 16.9691 | 3.47 | 492 | 4.887 | **<0.0001** |
| none - inter | 28.9838 | 4.25 | 492 | 6.816 | **<0.0001** |
| intra - inter | 12.0146 | 4.25 | 492 | 2.825 | **0.01** |
| exp_day = 14: | | | | | |
| none - intra | 17.6934 | 3.47 | 492 | 5.096 | **<0.0001** |
| none - inter | 27.8071 | 4.25 | 492 | 6.539 | **<0.0001** |
| intra - inter | 10.1137 | 4.25 | 492 | 2.378 | **0.046** |

| | | | | | |
|---|---|---|---|---|---|
| exp_day = 16: | | | | | |
| none - intra | 18.9492 | 3.47 | 492 | 5.457 | **<0.0001** |
| none - inter | 30.6419 | 4.25 | 492 | 7.206 | **<0.0001** |
| intra - inter | 11.6927 | 4.25 | 492 | 2.75 | **0.017** |
| **Cell shape after ~70 generations** | | | | | |
| contrast | estimate | SE | df | t-ratio | p-value |
| exp_day = 1: | | | | | |
| none - intra | -0.02821 | 0.00294 | 393 | -9.608 | **<0.0001** |
| none - inter | -0.02061 | 0.0036 | 393 | -5.733 | **<0.0001** |
| intra - inter | 0.007592 | 0.0036 | 393 | 2.111 | 0.09 |
| exp_day = 2: | | | | | |
| none - intra | -0.01818 | 0.00294 | 393 | -6.194 | **<0.0001** |
| none - inter | -0.02861 | 0.0036 | 393 | -7.957 | **<0.0001** |
| intra - inter | -0.01043 | 0.0036 | 393 | -2.9 | **0.011** |
| exp_day = 3: | | | | | |
| none - intra | -0.03177 | 0.00294 | 393 | -10.821 | **<0.0001** |
| none - inter | -0.03766 | 0.0036 | 393 | -10.474 | **<0.0001** |
| intra - inter | -0.00589 | 0.0036 | 393 | -1.638 | 0.23 |
| exp_day = 4: | | | | | |
| none - intra | -0.02409 | 0.00294 | 393 | -8.205 | **<0.0001** |
| none - inter | -0.02176 | 0.0036 | 393 | -6.051 | **<0.0001** |
| intra - inter | 0.002331 | 0.0036 | 393 | 0.648 | 0.79 |
| exp_day = 5: | | | | | |
| none - intra | -0.02041 | 0.00294 | 393 | -6.952 | **<0.0001** |
| none - inter | -0.01529 | 0.0036 | 393 | -4.252 | **0.0001** |
| intra - inter | 0.00512 | 0.0036 | 393 | 1.424 | 0.33 |
| exp_day = 6: | | | | | |
| none - intra | -0.02041 | 0.00294 | 393 | -6.952 | **<0.0001** |
| none - inter | -0.01157 | 0.0036 | 393 | -3.218 | **0.004** |
| intra - inter | 0.008839 | 0.0036 | 393 | 2.458 | **0.04** |
| exp_day = 8: | | | | | |
| none - intra | -0.01814 | 0.00294 | 393 | -6.179 | **<0.0001** |
| none - inter | -0.00442 | 0.0036 | 393 | -1.229 | 0.44 |
| intra - inter | 0.013722 | 0.0036 | 393 | 3.816 | **0.0005** |
| exp_day = 9: | | | | | |
| none - intra | -0.0128 | 0.00294 | 393 | -4.361 | **<0.0001** |
| none - inter | -0.00343 | 0.0036 | 393 | -0.955 | 0.61 |
| intra - inter | 0.009369 | 0.0036 | 393 | 2.606 | **0.026** |

| | | | | | |
|---|---|---|---|---|---|
| exp_day = 10: | | | | | |
| none - intra | -0.00695 | 0.0029 | 393 | -2.367 | **0.048** |
| none - inter | -0.00302 | 0.0036 | 393 | -0.841 | 0.68 |
| intra - inter | 0.00393 | 0.0036 | 393 | 1.092 | 0.52 |
| exp_day = 11: | | | | | |
| none - intra | -0.01023 | 0.0029 | 393 | -3.485 | **0.0016** |
| none - inter | -0.00991 | 0.0036 | 393 | -2.755 | **0.0169** |
| intra - inter | 0.00032 | 0.0036 | 393 | 0.09 | 0.9955 |
| exp_day = 12: | | | | | |
| none - intra | -0.00865 | 0.0029 | 393 | -2.945 | **0.0096** |
| none - inter | -0.00584 | 0.0036 | 393 | -1.624 | 0.24 |
| intra - inter | 0.00281 | 0.0036 | 393 | 0.781 | 0.72 |
| exp_day = 14: | | | | | |
| none - intra | -0.00984 | 0.0029 | 393 | -3.352 | **0.0025** |
| none - inter | -0.00393 | 0.0036 | 393 | -1.092 | 0.52 |
| intra - inter | 0.00591 | 0.0036 | 393 | 1.645 | 0.23 |
| exp_day = 16: | | | | | |
| none - intra | -0.0132 | 0.0029 | 393 | -4.498 | **<0.0001** |
| none - inter | -0.0036 | 0.0036 | 393 | -1.002 | 0.58 |
| intra - inter | 0.0096 | 0.0036 | 393 | 2.671 | **0.0214** |

**Table S4.** Scaling of *per capita* (per cell) photosynthesis and respiration rates with cell size for the three competition treatments together and for each competition treatment separately after ~70 generations of evolution with competitors. Irrespectively of how we analyse the data (together among competition treatments or separately), photosynthesis and respiration scale hyper-allometrically with cell size and decline over time as culture grow. We used linear models with $log_{10}$-transformed data.

| Rate | Competition | Slope estimate and CI for the effect of cell size | Slope estimate and CI for the effect of cell size × experiment day |
|---|---|---|---|
| *Photosynthesis* | All together | 1.31 (1.05; 1.58) | -0.06 (-0.09; -0.03) |
| | No competition | 1.35 (0.94; 1.75) | -0.07 (-0.12; -0.02) |
| | Intraspecific | 1.35 (0.91; 1.80) | -0.07 (-0.12; -0.009) |
| | Interspecific | 1.97 (1.36; 2.57) | -0.12 (-0.2; -0.05) |
| *Respiration* | All together | 1.28 (1.02; 1.55) | -0.05 (-0.08; -0.02) |
| | No competition | 1.34 (0.90; 1.79) | -0.06 (-0.12; -0.009) |
| | Intraspecific | 1.41 (0.98; 1.83) | -0.06 (-0.12; -0.007) |
| | Interspecific | 1.80 (1.23; 2.36) | -0.10 (-0.17; -0.03) |

**Table S5.** Coefficient estimates and post hoc tests for the effects of competition and initial cell density on max. growth rate ($r_{max}$, per day) and max. cell density ($K_{cells}$, cells µl$^{-1}$) of common garden experiments after ~35 (df = 42) and ~70 generations of evolution with competitors (df = 46). All tests use linear models, except for K at 35 generations for which we used generalized least squares with a treatment-specific variance because variances were heterogenous. None = competition-free. Intra = intraspecific competition. Inter = interspecific competition. P values < 0.05 are in bold.

| | | | | | |
|---|---|---|---|---|---|
| **~35 generations** | | | | | |
| **$r_{max}$** | | | | | |
| | Coefficient estimate | Std. Error | t-value | p-value | |
| Intercept | 568.28 | 14.70 | 38.67 | **< 0.0001** | |
| Initial density | 0.04 | 0.14 | 0.32 | 0.75 | |
| Competition (intra) | -71.85 | 20.73 | -3.47 | **0.001** | |
| Competition (inter) | -121.76 | 30.76 | -3.96 | **0.0003** | |
| contrast | estimate | SE | df | t-ratio | p-value |
| none - intra | 71.8 | 20.7 | 42 | 3.466 | **0.0035** |
| none - inter | 121.8 | 30.8 | 42 | 3.958 | **0.0008** |
| intra - inter | 49.9 | 30.9 | 42 | 1.614 | 0.25 |
| **$K_{cells}$** | | | | | |
| | Coefficient estimate | Std.Error | t-value | | p-value |
| Intercept | 2580.60 | 33.72 | 76.52 | | **< 0.0001** |
| Initial density | -0.56 | 0.62 | -0.90 | | 0.37 |
| Competition (intra) | 369.59 | 75.49 | 4.90 | | **< 0.0001** |
| Competition (inter) | 437.87 | 98.92 | 4.43 | | **0.0001** |
| contrast | estimate | SE | df | t-ratio | p-value |
| none - intra | -369.6 | 75.5 | 26.47 | -4.896 | **0.0001** |
| none - inter | -437.9 | 98.9 | 6.81 | -4.426 | **0.0080** |
| intra - inter | -68.3 | 116.0 | 11.73 | -0.589 | 0.8287 |
| **~70 generations** | | | | | |
| **$r_{max}$** | | | | | |
| | Coefficient estimate | Std. error | t-value | | p-value |
| Intercept | 464.36 | 16.78 | 27.67 | | **<0.0001** |
| Initial density | -0.005 | 0.07 | -0.07 | | 0.95 |
| Competition (intra) | 27.94 | 24.00 | 1.16 | | 0.25 |
| Competition (inter) | -28.88 | 29.05 | -0.99 | | 0.33 |
| **$K_{cells}$** | | | | | |
| | Coefficient estimate | Std. error | t-value | p-value | |
| Intercept | 2667.73 | 33.47 | 79.71 | **< 0.0001** | |
| Initial density | 0.14 | 0.14 | 0.96 | 0.34 | |
| Competition (intra) | 238.43 | 47.86 | 4.98 | **< 0.0001** | |
| Competition (inter) | 209.12 | 57.92 | 3.61 | **0.00075** | |
| contrast | estimate | SE | df | t-ratio | p-value |
| none - intra | -238.4 | 47.9 | 46 | -4.98 | **<0.0001** |
| none - inter | -209.1 | 57.9 | 46 | -3.61 | **0.002** |
| intra - inter | 29.3 | 58.2 | 46 | 0.50 | 0.87 |
